# Direct measurement of dynamic attractant gradients reveals breakdown of the Patlak-Keller-Segel chemotaxis model

**DOI:** 10.1101/2023.06.01.543315

**Authors:** Trung V. Phan, Henry H. Mattingly, Lam Vo, Jonathan S. Marvin, Loren L. Looger, Thierry Emonet

**Affiliations:** Department of Molecular, Cellular, and Developmental Biology, Yale University, New Haven, CT; Center for Computational Biology, Flatiron Institute, New York, NY; Janelia Research Campus, Howard Hughes Medical Institute, Ashburn, VA; Howard Hughes Medical Institute, Department of Neurosciences, University of California, San Diego, La Jolla, CA; Quantitative Biology Institute, Yale University, New Haven, CT; Department of Physics, Yale University, New Haven, CT

## Abstract

Chemotactic bacteria not only navigate chemical gradients, but also shape their environments by consuming and secreting attractants. Investigating how these processes influence the dynamics of bacterial populations has been challenging because of a lack of experimental methods for measuring spatial profiles of chemoattractants in real time. Here, we use a fluorescent sensor for aspartate to directly measure bacterially generated chemoattractant gradients during collective migration. Our measurements show that the standard Patlak-Keller-Segel model for collective chemotactic bacterial migration breaks down at high cell densities. To address this, we propose modifications to the model that consider the impact of cell density on bacterial chemotaxis and attractant consumption. With these changes, the model explains our experimental data across all cell densities, offering new insight into chemotactic dynamics. Our findings highlight the significance of considering cell density effects on bacterial behavior, and the potential for fluorescent metabolite sensors to shed light on the complex emergent dynamics of bacterial communities.

**SIGNIFICANCE STATEMENT:** During collective cellular processes, cells often dynamically shape and respond to their chemical environments. Our understanding of these processes is limited by the ability to measure these chemical profiles in real time. For example, the Patlak-Keller-Segel model has widely been used to describe collective chemotaxis towards self-generated gradients in various systems, albeit without direct verification. Here we used a biocompatible fluorescent protein sensor to directly observe attractant gradients created and chased by collectively-migrating bacteria. Doing so uncovered limitations of the standard chemotaxis model at high cell densities and allowed us to establish an improved model. Our work demonstrates the potential for fluorescent protein sensors to measure the spatiotemporal dynamics of chemical environments in cellular communities.

## INTRODUCTION

Motile bacteria, such as *Escherichia coli*, navigate complex and changing chemical signals (1) by performing a biased random walk (2): rotating its flagella counterclockwise propels the bacterium forward, until one or more motors reverse rotation direction, causing the cell to tumble and quickly reorient. By modulating their tumbling rate based on perceived changes in chemical concentration along their swimming trajectory (3,4), *E. coli* cells can climb gradients of attractants and descend gradients of repellents. This process, called chemotaxis, plays a critical role in bacterial pathogenesis (5), among numerous other phenomena.

Bacteria also actively modify their chemical environment and respond to these self-generated perturbations. For example, by consuming attractants and responding to the resulting gradients, populations of *E. coli* cells can migrate collectively (6–23). This process generates cell density waves such as traveling bands and rings that can migrate over large distances. In eukaryotes, such signaling molecule-induced collective cell migration events underlie cancer dynamics (24), immune system function (25), and development (26,27), among others.

The Patlak-Keller-Segel (PKS) model (28,29), and its extension to include growth and replication (12,15,18,19,30–33), is a standard mathematical framework for modeling chemotaxis, in particular collective migration, that has been widely used by researchers to investigate the interplay between bacterial collective behavior and chemical signaling (17,18,34,35,31,36–38,32,33). This standard chemotaxis model has proven to be a powerful tool for predicting emergent features observed in experiments, such as self-aggregation (9,21), pattern formation (39), and collective transport (10,21). However, the PKS model has never been directly verified, and more broadly our understanding of the intertwined dynamics of bacteria and their chemical environments is limited, due to the lack of techniques for measuring chemical concentration in real time.

Here we present a method to directly measure aspartate concentration with high spatial and temporal resolution, *in situ*, using a protein fluorescent -based sensor (40,41). With this sensor, we measure dynamic gradients generated by bacterial consumption during collective migration and test the validity of the standard PKS chemotaxis model. While the model agreed well with our data at low cell densities, it diverged at high cell densities. The inconsistencies between our experimental measurements and the PKS model can be resolved by introducing simple, biophysically-motivated modifications, in which both attractant consumption and bacterial chemotaxis are suppressed at higher cell densities. We expect that fluorescent protein-based sensors like the one used here will be valuable tools for accessing the spatiotemporal dynamics of chemical profiles in a variety of cellular systems.

## RESULTS

### Fluorescent sensor reveals spatiotemporal dynamics of bacteria-generated gradients during chemotaxis

To visualize self-generated gradients during collective bacterial migration, we repurposed a fluorescent sensor (iAspSnFR) for the attractant aspartate (Asp) (Fig. 1A), which some of us developed for use in neuronal systems (40,41). The sensor backbone is constructed from GltI, the periplasmic component of *E. coli*’s ABC transporter complex for import of the amino acids glutamate and aspartate (Asp). Two point mutations in the binding domain (S27A & S72P) make the sensor highly specific for Asp and insensitive to glutamate (SI Fig. S1). This ligand-binding domain is fused to circularly-permuted superfolder green fluorescent protein (GFP) to produce the fluorescent sensor. Upon binding to Asp, the sensor changes conformation, increasing its fluorescence emission intensity. We measured the calibration curve between fluorescence intensity and Asp concentration in microfluidic channels by mixing 1 μM of purified iAspSnFR protein with varying concentrations of Asp (Methods). These data fit an equilibrium binding model with a binding affinity of *K*_*d*_ = 20 ± 2 μM (Fig. 1B).

**Figure 1.**
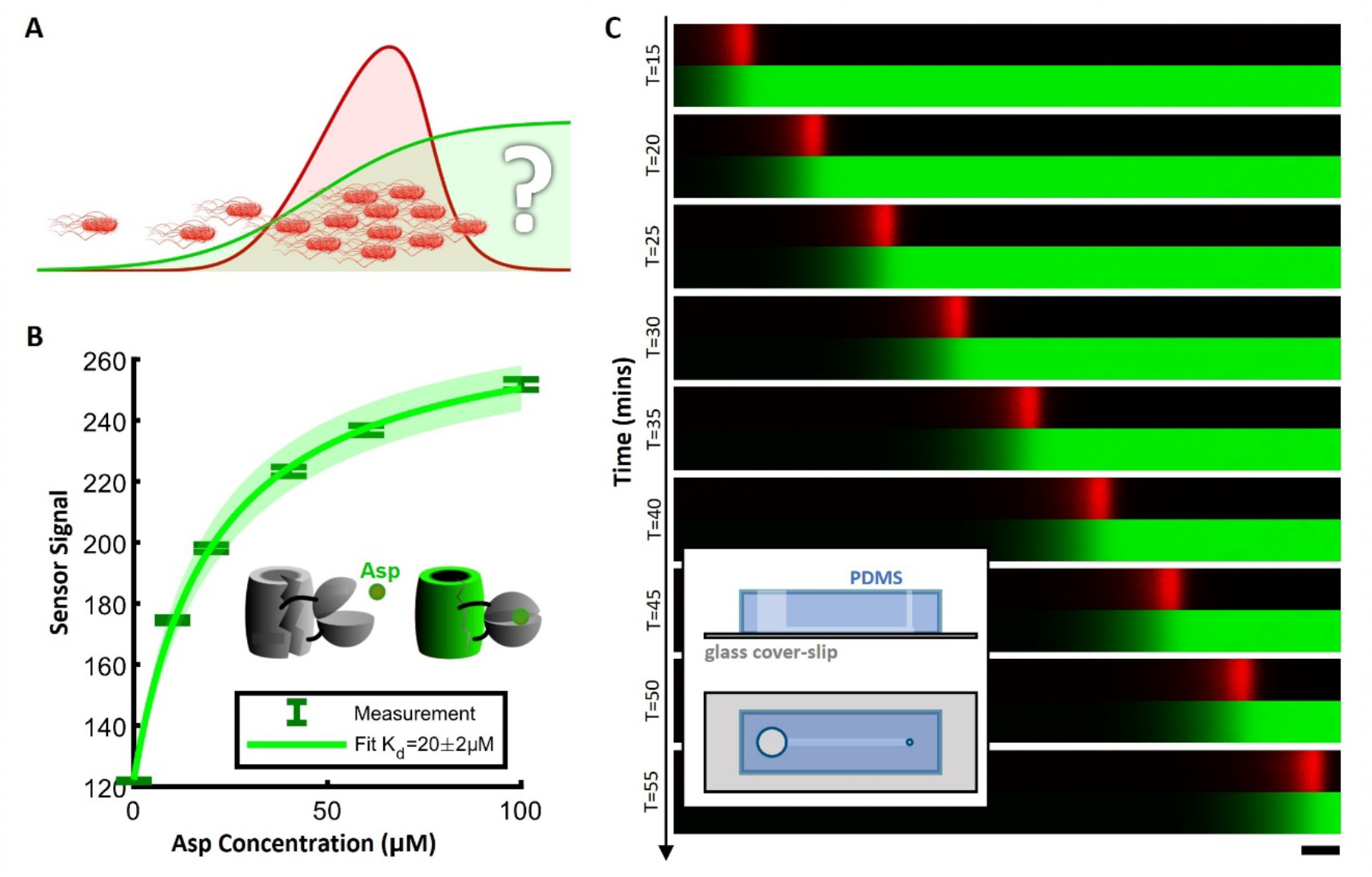
Fluorescent sensor reveals spatiotemporal dynamics of bacterially generated gradients during collective migration. **(A)** Bacteria (E. coli, red) respond to environmental chemical stimuli (attractant aspartate, green) that they themselves influence by consumption, enabling them to migrate collectively. **(B)** We repurposed a fluorescent sensor that changes conformation and increases fluorescence intensity after binding to aspartate (Asp). The plot shows the calibration curve from Asp concentration to sensor emission intensity. Data points are averages over an entire sweep of images along the microfluidic device. The same device was used for all Asp concentrations. Error bars are the standard deviation of pixel intensity over the entire sweep of images. Uncertainty of K_d_ is the 95% confidence interval of the fit. Shading is 95% confidence interval of the binding model’s predictions. **(C)** Time-lapse imaging of E. coli cells expressing the fluorescent protein mCherry2 (red) and the sensor (green) during collective migration in a microfluidic device (inset). Scale bar 1 mm. Inset: The microfluidic device consists of a long linear channel of length 20 mm, width 1.2 mm, and depth 100 μm, with two holes punched at the ends for filling in liquid medium (small hole) and inoculating bacteria (big hole).

Using the same microfluidic device and media as in the sensor calibration experiments, we generated collectively-migrating waves of red fluorescent protein (RFP)-labeled *E. coli* (Methods). Using 1 μM of sensor and 100 μM Asp ensured that the concentration of Asp in the wave was well within the range of sensitivity of the sensor and limited any potential effects of the sensor sequestering Asp from the bacteria. With this experimental setup, we used time-lapse imaging to simultaneously measure the spatial density of bacteria and the dynamic profile of Asp (Fig. 1C; Methods). By separately measuring RFP fluorescence intensity for several known densities of bacteria, we interpolated the absolute density of bacteria in the traveling waves, in units of optical density (OD) at 600 nm. To the best of our knowledge, this is the first time that both the concentrations of bacteria and attractant have been directly measured with high spatial resolution in real time.

### The standard model for collective chemotactic migration describes the bacterial wave at low cell density but breaks down at high cell density

Numerous theoretical and experimental studies have used variations of the classic PKS model to describe collective chemotactic migration (28,29). This model consists of two coupled partial differential equations (PDEs) that describe the dynamics of Asp concentration 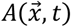 and bacterial density 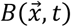 The Asp dynamics include diffusion and consumption by the bacteria:

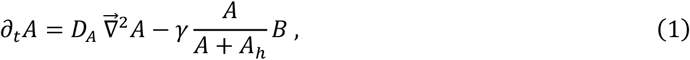

where *D*_*A*_ = 800 μm^2^/s is the diffusion coefficient of Asp in water (17,18,42–44), *γ* is the maximum consumption rate, and *A*_*h*_ is the Asp concentration at which bacteria consume Asp at half-maximal rate (17,18,45). The bacterial dynamics include cell diffusion, chemotaxis, and growth:

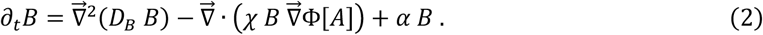

Here, 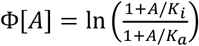 is the cells’ perceived signal, where *K*_*i*_ ∼ 1 μM (46–48) and *K*_*a*_ ≫ *A* are the receptor-ligand dissociation constants when receptors are in the inactive and active states, respectively (49). *D*_*B*_ and *χ* are the diffusion and chemotaxis coefficients of the bacteria, and *α* is the growth rate, which we measured to be *α* = 0.38 ± 0.02/h under our experimental conditions (Methods).

With the ability to directly measure both bacterial density and *in situ* Asp concentration, we could assess the PKS model of collective bacterial migration. During collective migration along the linear channel in our device, the profiles of both the bacterial density *B* and Asp concentration *A* reached a quasi-steady state. That is, the profiles essentially stopped changing in the frame of reference moving with the bacterial wave, defined by *z* = *x* − *ct*, where *c* is the speed of the traveling wave, and *z* = 0 is defined as the peak of bacterial density. Converting Eqs. (1) and (2) to the co-moving frame and rearranging terms so that all unknown parameters are on the right-hand side (RHS), we arrive at:

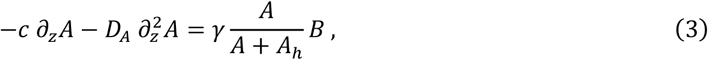

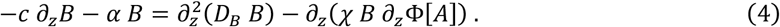

The consumption parameters *γ* and *A*_*h*_, although measured before under other conditions (17,18), can vary with bacterial strain, growth media, temperature, experimental media, *etc*. - as can *χ* and *D*_*B*_. Therefore, to test the ability of this model to quantitatively capture our data, we fit these unknown model parameters by computing the left-hand side (LHS) of Eqs. (3) and (4) from the data and then minimizing the sum of squared deviations with the right-hand side (RHS) (Methods).

These PDEs fit the data well for low cell density bacterial waves (peak cell density OD = 0.7) with *γ* = 0.45 ± 0.04 μM/OD/s, *A*_*h*_ = 7.6 ± 3.7 μM and *D*_*B*_ = 400 ± 100 μm^2^/s, *χ* = 3300 ± 230 μm^2^/s (Fig. 2A, B, C). The fit values are in reasonable ranges, compared to the literature (17,18,50–52). However, the model predictions with the same parameters values clearly deviated from the data in high cell density waves (peak cell density OD = 5.1) (Fig. 2D, E, F). For model predictions on intermediate-density waves, see SI Fig. S2. Our findings reveal that the standard model for chemotaxis fails to accurately describe the behavior of high-density bacterial populations.

**Figure 2.**
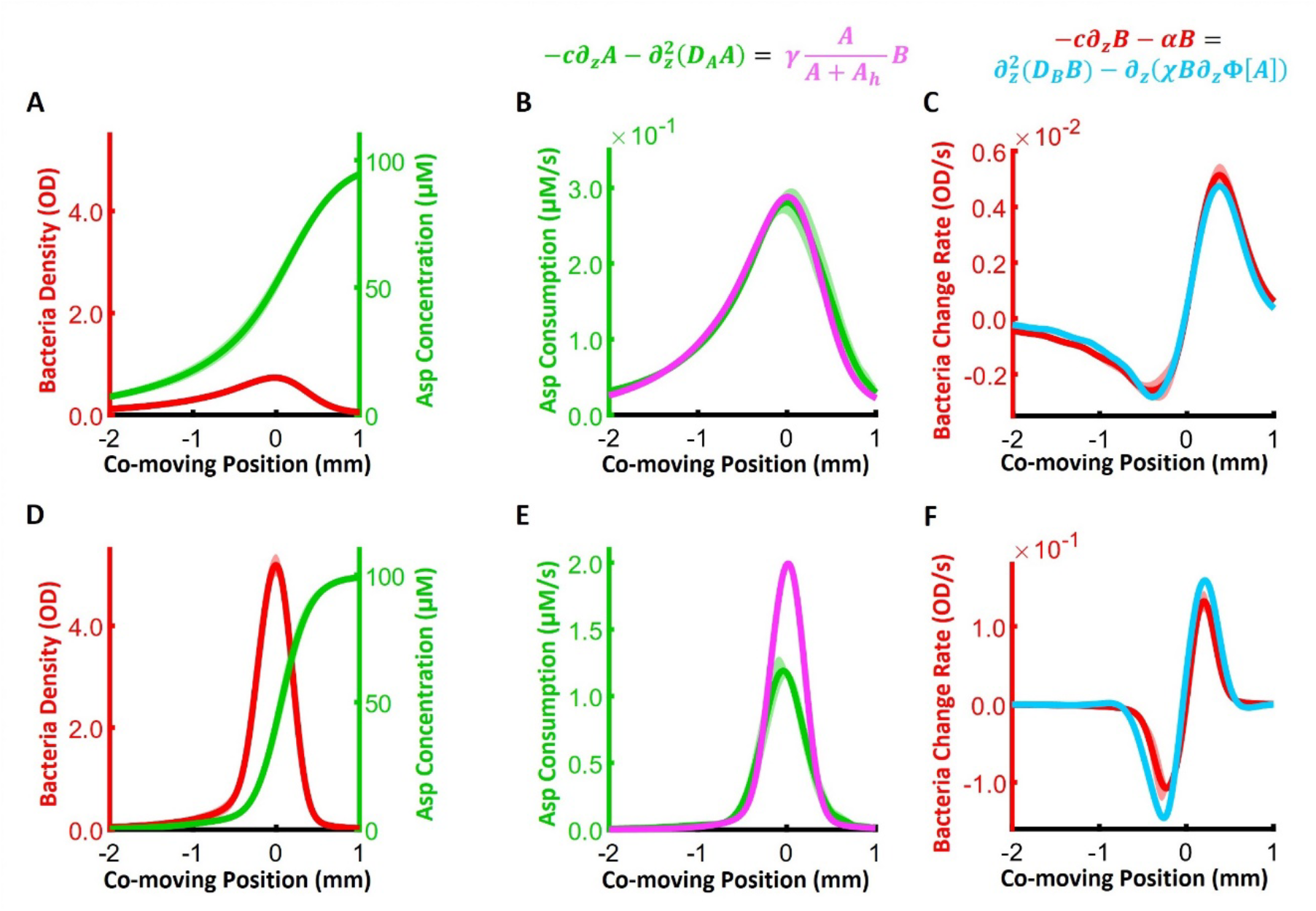
The standard model for chemotaxis, the Patlak-Keller-Segel (PKS) model, describes the bacterial wave at low cell density but breaks down at high density. **(A)** The quasi-steady state profiles in the co-moving frame, of bacteria density (red) and attractant Asp concentration (green), during low cell density migration (OD = 0.7). **(B)** Fitting the Asp dynamics of the PKS model to experimental data (above). A and B are the Asp concentration and the bacteria density respectively, c is the wave speed, D_B_ is the molecular diffusion of Asp, γ is the maximum consumption rate, A_h_ is the half-max of the consumption rate, and z = x − ct is the coordinate in the co-moving frame (z = 0 is defined as the peak of bacterial density). Green: LHS of the Asp dynamics equation with A and c = 4.4 ± 0.1 μm/s measured and D_B_ = 800 μm^2^/s (17,18,42–44). Throughout, uncertainty of c is the standard error of the mean. Magenta: RHS of the Asp dynamics equation with B measured, and γ = 0.45 ± 0.04 μM/OD/s and A_h_ = 7.6 ± 3.7 μM fit to match the LHS. **(C)** Fitting the bacteria dynamics of the PKS model to experimental data (above). 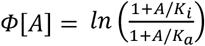 is the cells’ perceived signal (K ≫ A and K = 1 μM (46–48)), D is the effective bacterial diffusivity, χ is the chemotactic coefficient, and α = 0.4/hr is the measured growth rate (Methods). Red: LHS of the bacteria dynamics equation with α, c and B measured. Cyan: RHS of the bacterial dynamics equation with D_B_ = 400 ± 100 μm^2^/s and χ = 3300 ± 230 μm^2^/s fit to match the LHS. **(D**,**E**,**F)** is the same as **(A**,**B**,**C)** for a high cell density wave. **(E**,**F)** magenta and cyan lines are prediction using the same parameter values for γ, A_h_, D_B_, χ as in **(B**,**C)**. Here c = 7.7 ± 0.2 μm/s. The shading represents the standard deviation across replicates. N = 6 replicates for **(A**,**B**,**C)** and N = 4 for **(D**,**E**,**F)**.

### Attractant dynamics are consistent with cell density-dependent consumption rate

At high cell densities, the model fails to describe both Asp and bacterial dynamics. The predictions of Asp dynamics break down because they overestimate the rate at which bacteria consume the attractant near *z* = 0, where cell density is highest (Fig. 2E). This disagreement gets worse as bacterial density increases (SI Fig. S2HK). Therefore, we propose the following phenomenological correction to the maximum consumption rate, in which the bacteria consume aspartate more slowly at higher local cell density:

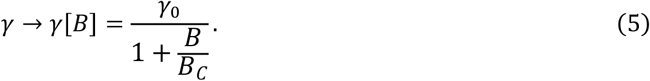

For the parameter values *γ*_0._ = 0.47 ± 0.06 μM/OD/s, *B*_C_ = 5.8 ± 2.0 OD and *A*_*h*_ = 3.7 ± 3.2 μM, the RHS is in reasonable agreement with the LHS of Eq. (3), for the entire range of wave cell densities observed (Fig. 3). The fit values for *γ*_0._ and *A*_*h*_ are also reasonable compared to the literature (17,18).

**Figure 3.**
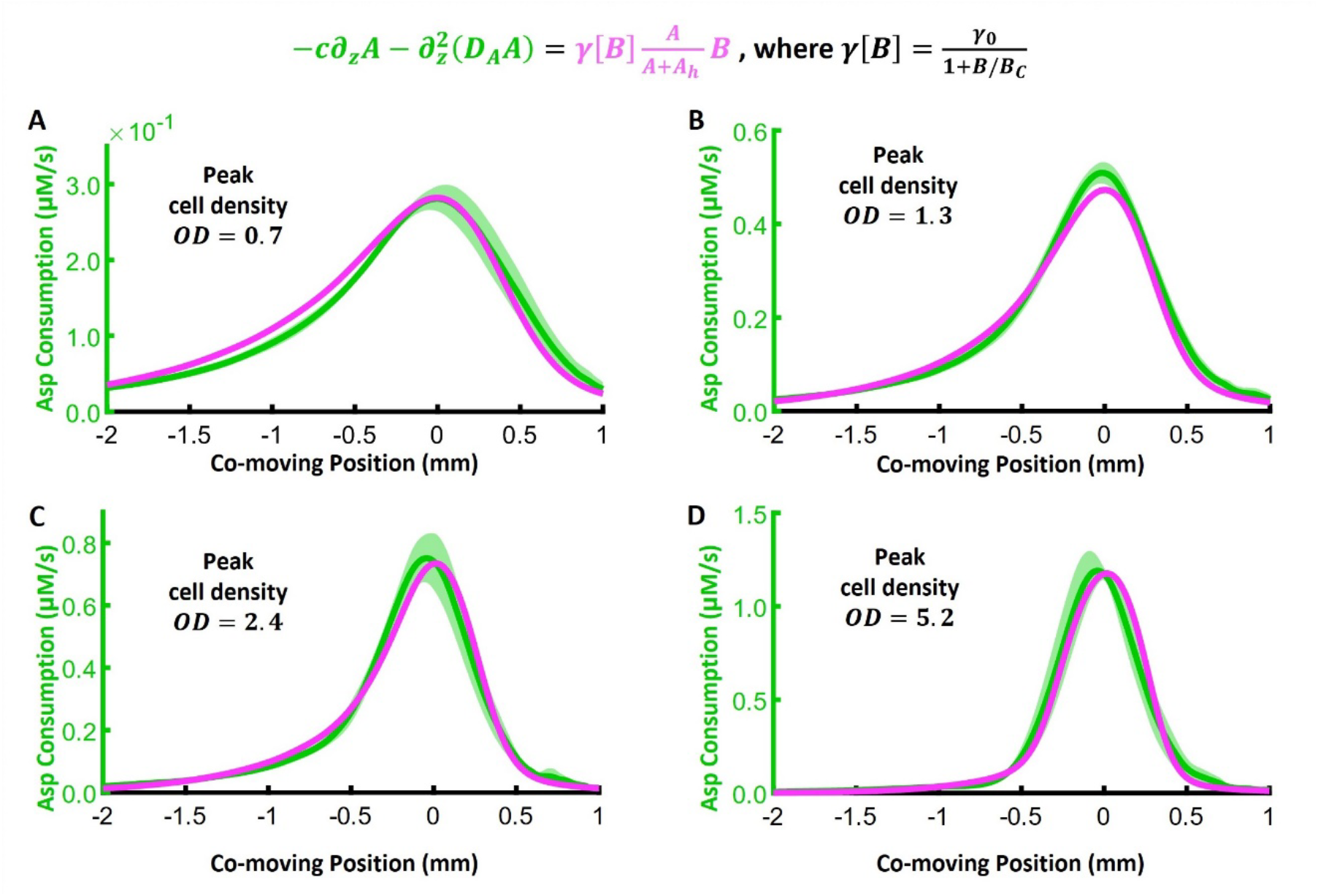
Asp dynamics are consistent with cell density-dependent consumption rate. Top: modified Asp consumption rate. Green: Asp consumption rate (LHS) calculated from the data. Magenta: Asp consumption rate (RHS) with parameters γ_0._ = 0.47 ± 0.06 μM/OD/s, B_C_ = 5.8 ± 2.0 OD and A_h_ = 3.7 ± 3.2 μM fit to match the LHS. These are shown for waves with peak bacterial cell density of OD = 0.7 **(A)**, 1.3 **(B)**, 2.4 **(C)**, and 5.2 **(D)**. The parameters were fit to all data simultaneously. The shading represents the standard deviation across replicates: N = 6 for (A), N = 9 for (B), N = 5 for (C), and N = 4 for (D).

One plausible mechanism for a cell density-dependent consumption rate of Asp is spatial variations in the concentration of dissolved oxygen, which is required by *E. coli* to consume Asp (17,53). Oxygen is consumed by *E. coli* (6,17), but it can be replenished by permeation through the PDMS chip (54). In microfluidic devices where the channel depth is much smaller than the width of the traveling wave, these two effects can balance out such that the local drop in oxygen concentration is directly proportional to the local cell density (see SI for derivation). Indeed, in a similar microfluidic system, oxygen levels in traveling waves of bacteria were shown to be lower in regions of higher cell density (17).

### The shape of the attractant gradient reveals a reduction of the chemotactic coefficient at high cell density

The standard PKS model prediction for the flux of bacteria also deviates from our observations in waves with high cell density. Like the Asp consumption dynamics, this disagreement between theory and experiment increases with cell density. The deviation is most notable behind *z* = 0 (Fig. 2F), *i*.*e*., the region behind the peak of cell density, where the bacterial flux is dominated by chemotaxis (29,32). Therefore, we inspected the profile of perceived signal Φ[*A*], whose gradient drives chemotactic waves. As the maximum cell density in the wave increased (Fig. 4A), the perceived signal Φ[*A*] exhibited regions with two different slopes: a steeper slope near the peak of cell density, and a shallower slope behind it (Fig. 4B). These two regions were even more apparent when we plotted the gradient of perceived signal Φ[*A*] (Fig. 4C), which showed a distinct peak close to the peak of cell density that increased with maximum cell density.

**Figure 4.**
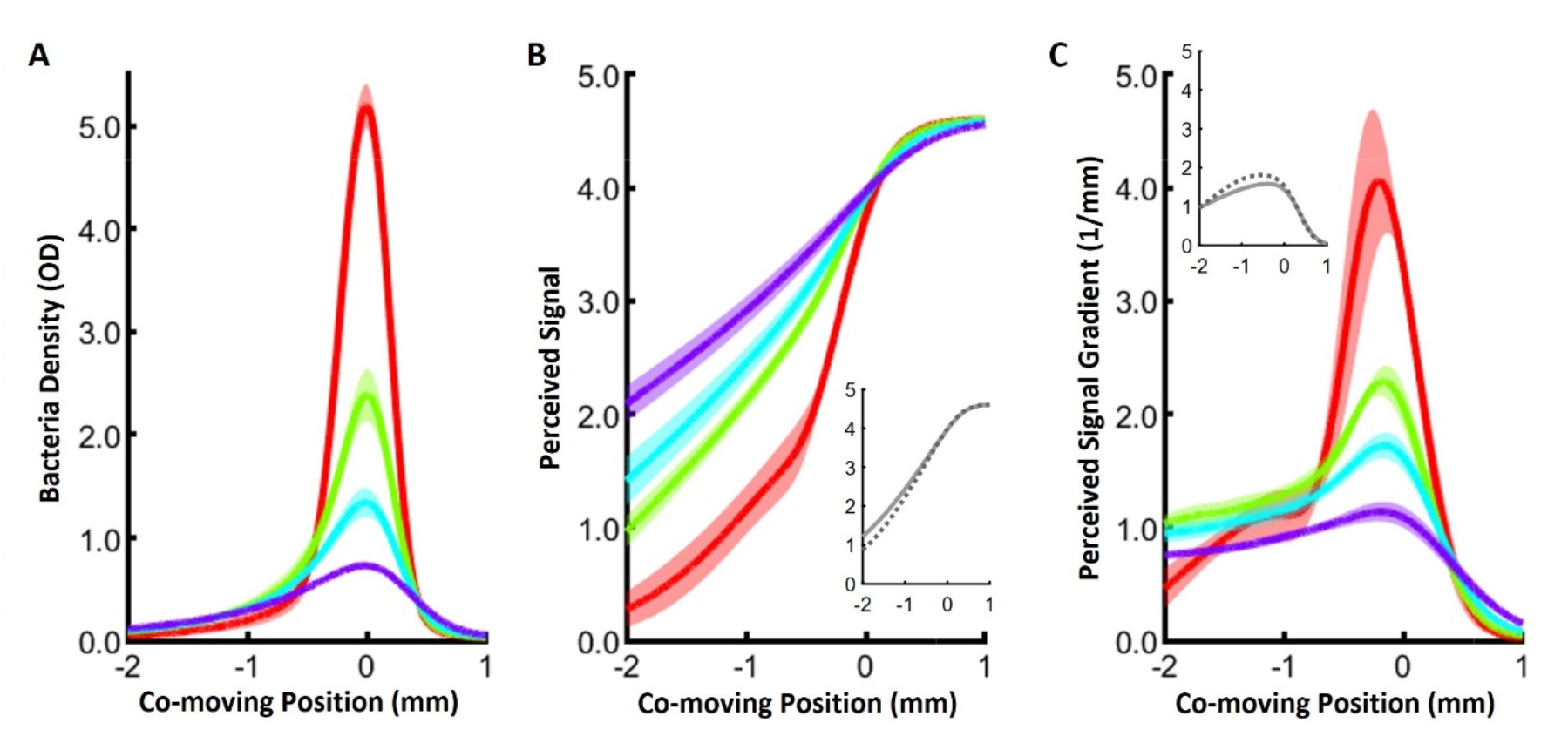
The gradient of the measured perceived signal cannot be explained by the standard model for chemotaxis. **(A)** Cell density profile for four waves with increasing peak cell density: OD = 0.7 (purple, N = 6), 1.3 (cyan, N = 9), 2.4 (green, N = 5), and 5.2 (red, N = 4). Shade is standard deviation across replicates. **(B)** Corresponding perceived signal 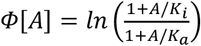 exhibits a sudden change of slope at the back of the wave that becomes more prominent with higher cell density. Inset: Simulations of the PKS model using the proposed cell density-dependent Asp consumption rate but a constant chemotactic coefficient χ show no sharp change in the slope of the perceived signal. Simulations with (full line) and without (dashed line) the contribution of phenotypic diversity. **(C)** Corresponding gradient of the perceived signal ∂_z_Ф[A] shows a distinct peak that becomes more prominent for high cell density waves. Inset: plots from the PKS simulations as in **(B)** shows that PKS equations cannot explain this feature even with the effects of phenotypic diversity (17) included.

This drastic change in the slope of the attractant gradient arguably provides the clearest evidence for a breakdown of the standard PKS model. The speed of the wave *c* is set by how fast the cells deplete the attractant (29). Then to travel at that speed, the local gradient steepness ∂_*z*_Φ[*A*] must be related to the chemotactic coefficient of the traveling cells according to ∂_*z*_Φ[*A*] ∼ *c*/*χ*(*z*) (29,32,33). The two slopes indicated that the bacteria dynamics cannot be explained by a single value of the chemotactic coefficient. Simulations confirmed that two distinct slopes do not emerge in the PKS model, even after including our modification to the Asp consumption rate in Eq. (5) (Fig. 4BC insets).

Phenotypic diversity is known to cause individual cells to have different chemotactic coefficients (55,56). Since the gradient is shallow at the front of the wave and steeper towards the back, high-performing phenotypes (high-*χ* cells) localize to the front, and low-performing phenotypes localize further back, equalizing migration speed throughout the wave (17). This arrangement is stable to perturbations in the region where the perceived gradient monotonically increases with position *z* from front to back: when a cell moves away from its stable point in the wave, the steeper gradient behind that point makes the cell speed up, while the shallower gradient ahead makes it slow down, returning the cell to the stable point in both cases.

But this arrangement cannot explain why cells are still able to travel together in the region of the wave where the gradient is decreasing towards the back, which becomes pronounced in the high cell density case. In this region, the cells’ chemotactic coefficients must increase instead of decrease towards the back, which should not be stable according to the logic described above. Simulations of the PKS model with and without phenotypic diversity confirmed that spatial sorting alone could not explain the emergence of a peak in the signal gradient at high cell density (Fig. 4BC insets). Therefore, there must be another, unknown mechanism that maintains the stability of an arrangement in which some cells with high *χ* are located behind cells with low *χ*_0_, particularly in waves with high cell density.

Stability requires that this mechanism be a local, dynamic effect on top of cells’ intrinsic chemotactic coefficients, *χ*_0_. In particular, we propose a phenomenological model in which cells’ chemotactic coefficients are reduced in regions of high cell density:

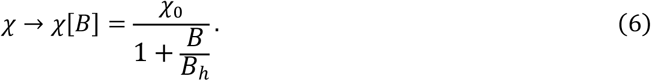

In this model, *χ*_0._ can vary among cells, but it is divided by a cell density-dependent factor. When a cell in the high-density region falls behind, although the gradient gets shallower, its dynamic *χ*_0_ increases according to Eq. (6), and the net effect is to return the cell to the high-density region. Fitting the RHS to the LHS of Eq. (4) using this correction showed good agreement with the data for all cell densities observed (Fig. 5), with parameter values *D*_*B*_ = 440 ± 90 μm^2^/s, *χ*. = 3800 ± 450 μm^2^/s, and *B*_*h*_ = 7.7 ± 3.1 OD.

**Figure 5.**
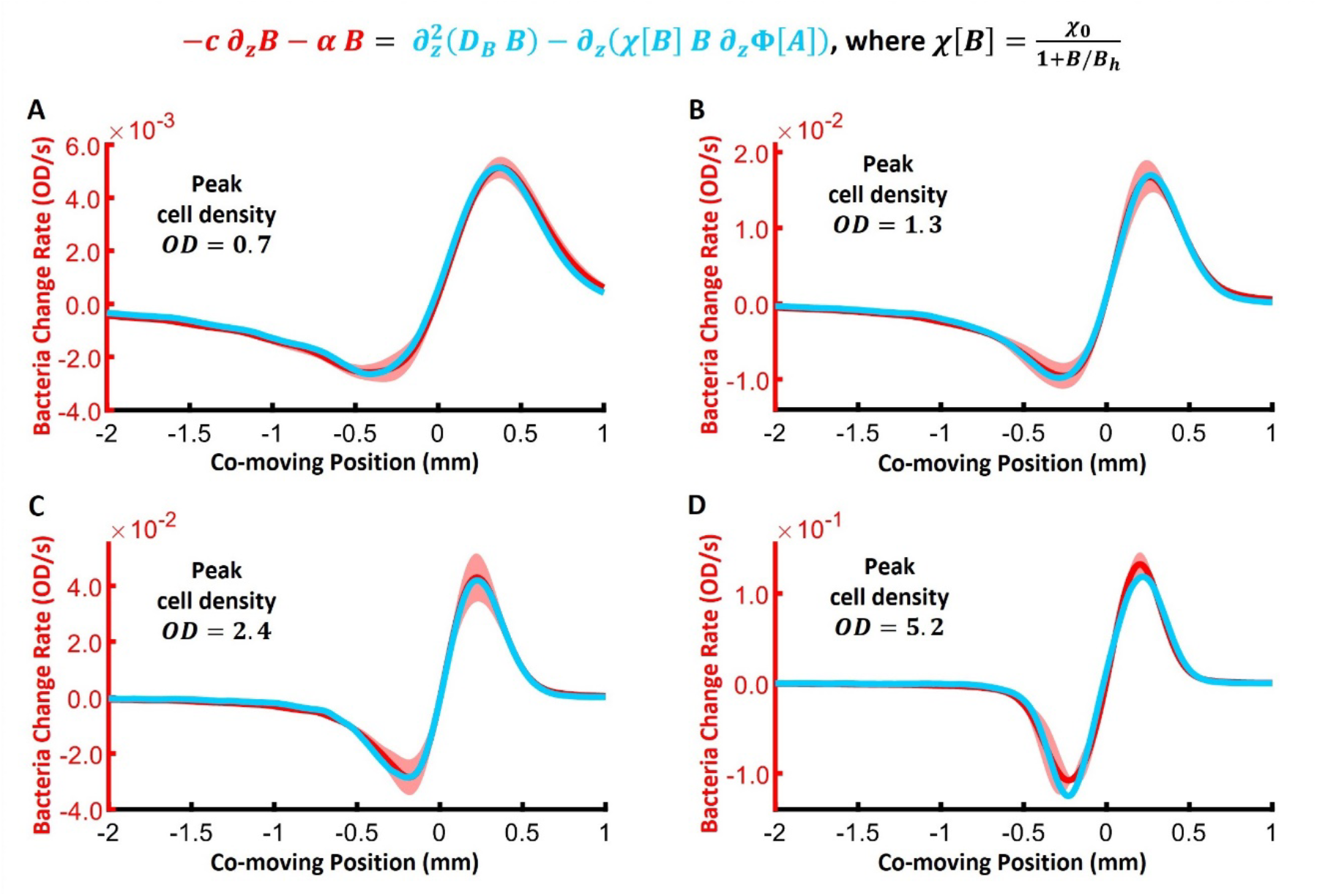
Bacterial dynamics are consistent with a reduction of the chemotactic coefficient when cell density becomes large. Top: modified chemotactic coefficient. Red: rate of change of cell density (minus growth) (LHS) calculated from the data. Cyan: spatial derivative (divergence) of the flux of cell density (RHS) with parameters D_B_ = 440 ± 90 μm^2^/s, χ. = 3800 ± 450 μm^2^/s, B_h_ = 7.7 ± 3.1 OD fit to match the LHS. These are shown for waves with peak bacterial cell densities of OD = 0.7 **(A)**, 1.3 **(B)**, 2.4 **(C)**, and 5.2 **(D)**. The parameters were fit to all data simultaneously.

We note that chemotaxis to oxygen can also affect group migration of *E. coli* (6). But chemotaxis to oxygen would produce the opposite of the observed effect: an oxygen gradient would point forward at the front and backward at the back, since the local drop in oxygen is proportional to the local cell density (SI). With an additional chemotactic signal like this, coherent migration would require that the Asp gradient be shallower at the front and steeper at the back, contrary to our experimental observations. Thus, chemotaxis to oxygen gradients cannot explain the shape of the measured gradient.

Reduction of the chemotactic coefficient with increasing cell density has been reported before in experiments measuring the drift velocity of individual cells in a fixed gradient of the non-metabolizable attractant methyl-aspartate (57). In the theory of that paper, *χ*[*B*] has almost the same form as Eq. (6), except with an additional, higher-order *B*^2^ term in the denominator. The cell density *B*_*h*_ at which this effect becomes significant in our data is similar to the value reported there. When we fit the full functional form, we found that the coefficient multiplying the quadratic term was near zero, therefore we did not include that term. Comparing agent-based simulations to experiments, those authors found that hydrodynamic interactions among cells increased their effective rotational diffusion at high cell densities, thus impairing their ability to bias their swimming direction in response to chemical signals. Here, we find that this cell density-dependent reduction in the chemotactic coefficient steepens the attractant gradient in the region where cell density is highest, compensating for the reduction in chemotactic coefficient there.

## DISCUSSION

By repurposing a fluorescent sensor for the amino acid aspartate, here we have directly measured dynamic chemoattractant gradients generated by bacterial cells, with high spatial and temporal resolution. Using these measurements, we tested a classic and extensively-used model of collective chemotactic migration, the Patlak-Keller-Segel model (28,29), extended to include growth (12,15,18,19,30–33). We discovered that while the standard PKS model provides a satisfactory description of travelling waves at low cell density, it fails to explain experimental observations at high cell densities. Specifically, aspartate consumption is lower than predicted at high densities, and the perceived signal behind the peak of the wave exhibits two increasingly distinct slopes. We find that phenomenological extensions to the PKS model, in which aspartate consumption and chemotaxis performance decrease with increasing cell density, explain the data over a wide range of cell densities. Our findings imply that the impacts of cell density on bacterial collective behavior are profound, even when the cell volume fraction is small (<1%).

Our understanding of many cell-biological processes, such as wound healing (58,59), development (60), and biofilm formation (61–64), is likely limited by the difficulties of measuring the spatiotemporal dynamics of chemical concentrations experienced by and shaped by cells. Julius Adler performed the first measurements of cell density and attractant concentrations in traveling waves of bacteria (6), but they were limited to a spatial resolution of a centimeter and temporal resolution of an hour. Here by contrast, we measured the dynamics of aspartate concentration with micron and minute resolution, with the latter limited by the time needed to take sweeps of images over a large field of view. Common approaches typically involve fixation (65,66), which can provide spatial structure and snapshots, but with limited dynamic information – and frequent damage to the processes under study. Other alternatives include invasive probes (67) or small molecule dyes that are toxic to cells (68). Biocompatible, protein-based fluorescent sensors like the one used here provide a powerful method for addressing these difficulties.

## METHODS

### Microfluidic device design and fabrication

Microfluidic devices were constructed from the biocompatible and oxygen-permeable silicone polymer polydimethylsiloxane (PDMS) on cover slips following standard soft lithography protocols. The master mold for the device is a silicon wafer featuring a long linear channel of width 1.2 mm, length 2 cm, and height 100 μm, created using ultraviolet (UV) photoresist lithography with SU-8 negative resist (SU8 3010, Microchem). The resists were then cured using UV light exposure through photomasks designed in CAD software and printed by CAD/Art Services, Inc. (Bandon, Oregon), again following photoresist manufacturer specifications. Subsequently, the wafer was baked, and the uncured photoresist was dissolved. After curing the SU-8 coat, the features were baked further, and after completion a protective coat of silane was applied by vapor deposition.

To cast the device, the wafer was coated with a 5-mm layer of degassed 10:1 PDMS-to-curing agent (Sylgard 184, Dow Corning). The layer was baked overnight at 70°C and allowed to cool. Then the devices were cut out, and holes of diameter size 2 mm and 0.25 mm were punched at the two ends of the long linear channel, making external connections with the channel from the outside. The PDMS devices were cleaned with transparent adhesive tape (Magic Tape, Scotch) followed by rinsing with (in order) isopropanol, methanol, and Millipore filtered water, air-drying between each rinse. The glass was rinsed the same way, with acetone, isopropanol, methanol, and Millipore-filtered water. The PDMS was tape-cleaned an additional time, and then the devices and 24 mm *×* 50 mm glass coverslips (#1.5) were placed in a plasma treatment oven (Harrick Plasma) under vacuum for 60 s. The PDMS was then laminated to the cover slip and baked on a hotplate at 80°C for 15 mins to establish a covalent bond. Devices were stored at room temperature and used within 12 h.

### Protein purification

The Asp sensor used here, iAspSnFR (manuscript in preparation), is a precursor of iGluSnFR3 (69), with additional mutations to the binding pocket, S27A and S72P. Plasmids carrying the sensor were transformed into *E. coli* BL21(DE3) cells (lacking pLysS). Protein expression was induced by growth in liquid auto-induction medium supplemented with 100 μg/mL ampicillin at 30°C (70). Proteins were purified by immobilized Ni-NTA affinity chromatography (71) in 0.1 M sodium phosphate buffer containing 1 M NaCl, pH 7.4. The sensor protein was eluted with a 120-mL gradient from 0 mM to 200 mM imidazole. Final concentrations of sensor were typically ∼100 μM.

### Strains, growth conditions, and sample preparation

The bacterial strain used in this study is HE205 (18), a motile variant of *E. coli* K-12 strain NCM3722 whose physiology has been well-characterized (72–78). HE205 was transformed with plasmid pZA31 carrying genes for the red fluorescent protein mCherry2 (79) and chloramphenicol antibiotic resistance, both under the constitutive promoter pTet. We grew cells in MOPS glycerol medium, made from 100 mL/L 10x MOPS N-C-, with additional components added in the following order to avoid precipitation of salts: 83.7 g/L MOPS adjusted to pH 7.0 with KOH, 7.12 g/L tricine adjusted to pH 7.0 with KOH, 0.0278 g/L FeSO_4_ 7H_2_O, 0.481 g/L K_2_SO_4_, 0.000555 g/L CaCl_2_, 1.06 g/L MgCl_2·_6H_2_O, 29.1 g/L NaCl, 0.230 g/L K_2_PO_4_, 20 mM NH_4_Cl, 4 mL/L glycerol, then sterile filtered. The medium used in the travelling wave experiments is the same as the growth medium, but with Asp added to a final concentration of 100 μM and iAspSnFR added to a final concentration 1 μM.

Cells first grew overnight, then were diluted 100x and regrown until mid-exponential phase OD600 = 0.25. After that, cells were washed twice in the growth medium and resuspended in the experimental medium at cell densities (OD600) of 0.1, 0.3, 3.0 and 6.0 to generate waves of different cell densities. After filling the microfluidic device with experimental medium from the smaller hole, that hole was sealed with a piece of transparent adhesive tape (Magic Tape, Scotch) to prevent flow during the experiment caused by differences in hydrostatic pressure across the device. Cells were then inoculated in the larger hole, and a drop of mineral oil was also added to prevent evaporation.

### Imaging and analysis

Migrating waves were imaged on an inverted microscope (Nikon Eclipse Ti-E) equipped with a custom environmental chamber (50% humidity and 30°C). A custom MATLAB script was used to control the microscope and its automated stage (Prior) via the MicroManager interface (80). Time-lapse images (fluorescence: RFP and GFP) of the bacteria and the Asp sensor were acquired using a Hamamatsu ORCA-Flash4.0 V2 camera (2048 *×* 2048 array of 6.5 μm *×* 6.5 μm pixels), a 10x objective (Nikon CFI Plan Fluor, N.A. 0.30, W.D. 16.0 mm) and an LED illuminator (Lumencor SOLA light engine, Beaverton, OR). Starting at the origin (closed beginning of the long linear channel) with a specified color channel, the motorized stage moved along the channel and paused every 1/3 the width of one frame to take fluorescence images (exposure time 122 ms for both channels). All images in the mCherry2 (RFP) channel were taken first, followed by images in the iAspSnFR (GFP) channel. Image sweeps were taken every 5 mins.

Raw RFP and GFP images from each sweep were first stitched together into “frames”. Then each frame was normalized by the first GFP frame, which was taken before the cells entered the channel, to eliminate the effects of variations in the height of the channel on fluorescence intensity. The normalized frames were then rotated so that the channel was aligned with horizontal axis, and pixels outside of the channel were cropped out. Finally, the fluorescence intensity inside the channel was averaged over the width the channel, *i*.*e*., the direction perpendicular to the direction of motion of the wave. Spatial derivatives were computed by approximating the measured profiles with smoothing spline functions using the MATLAB function *fit* with the ‘smoothingspline’ option and computing the derivative of the splines. Asp concentration profiles and cell density profiles were averaged over replicates with similar peak cell density. Models were fit to data from all cell density conditions simultaneously by minimizing the sum of squared deviations between the data and the model, normalized by the norm of the data function, using the MATLAB function *lsqnonlin*, specifically in the 3mm region around the cell density peak.

### Growth rate measurement

Cells were grown overnight in the experimental medium (without Asp), then diluted and grown to OD 0.25, as above (Asp has a small effect on the growth rate in this medium (18)). Then, cells were inoculated in the microfluidic device, as above. The device with cells was loaded onto the microscope with the environmental chamber and imaged by taking sweeps of the RFP channel every 5 mins for 6 hours. Images were analyzed as above to extract the average RFP intensity, which was converted to OD. The growth rate was estimated by fitting a line to the logarithm of OD versus time during exponential growth.

### Simulation

Simulations were performed using the same custom MATLAB code reported recently (33), with the Asp consumption rate modified as in Eq. (5).

## Code availability

Simulation code is available at https://github.com/emonetlab/ks.

## Data availability

The experimental data will be available on datadryad.org upon publication. The iAspSnFR sensor is available upon request from Jonathan Marvin.

## ACKNOWLEDGEMENTS

We thank Thomas Lamot for extensive help setting up the experimental assay. We also thank Jeremy Moore, Keita Kamino, Jyot Antani, and Truong H. Cai for helpful discussions. This work was supported by NIH awards R01GR110270 (T.E.) and F32GM131583 (H.H.M.).

## Notes

### Competing Interest Statement

The authors have declared no competing interest.

## REFERENCES

1. Sourjik V, Wingreen NS. Responding to chemical gradients: bacterial chemotaxis. Current Opinion in Cell Biology. 2012 Apr 1;24(2):262–8.

2. Berg HC. E. coli in motion. New York: Springer; 2004. 133 p. (Biological and medical physics series).

3. Segall JE, Block SM, Berg HC. Temporal comparisons in bacterial chemotaxis. PNAS. 1986 Dec 1;83(23):8987–91.

4. Parkinson JS, Hazelbauer GL, Falke JJ. Signaling and sensory adaptation in Escherichia coli chemoreceptors: 2015 update. Trends in Microbiology. 2015 May 1;23(5):257–66.

5. Zhou B, Szymanski CM, Baylink A. Bacterial chemotaxis in human diseases. Trends in Microbiology. 2023 May 1;31(5):453–67.

6. Adler J. Chemotaxis in Bacteria. Science. 1966 Aug 12;153(3737):708–16.

7. Holz M, Chen SH. Quasi-elastic light scattering from migrating chemotactic bands of Escherichia coli. Biophysical Journal. 1978 Jul 1;23(1):15–31.

8. Wolfe AJ, Berg HC. Migration of bacteria in semisolid agar. PNAS. 1989 Sep 1;86(18):6973–7.

9. Park S, Wolanin PM, Yuzbashyan EA, Lin H, Darnton NC, Stock JB, et al. Influence of topology on bacterial social interaction. PNAS. 2003 Nov 25;100(24):13910–5.

10. Lambert G, Liao D, Austin RH. Collective Escape of Chemotactic Swimmers through Microscopic Ratchets. Phys Rev Lett. 2010 Apr 22;104(16):168102.

11. Saragosti J, Calvez V, Bournaveas N, Perthame B, Buguin A, Silberzan P. Directional persistence of chemotactic bacteria in a traveling concentration wave. PNAS. 2011 Sep 27;108(39):16235–40.

12. Koster DA, Mayo A, Bren A, Alon U. Surface Growth of a Motile Bacterial Population Resembles Growth in a Chemostat. Journal of Molecular Biology. 2012 Dec 7;424(3):180–91.

13. Emako C, Gayrard C, Buguin A, Almeida LN de, Vauchelet N. Traveling Pulses for a Two-Species Chemotaxis Model. PLOS Computational Biology. 2016 Apr 12;12(4):e1004843.

14. Ni B, Ghosh B, Paldy FS, Colin R, Heimerl T, Sourjik V. Evolutionary Remodeling of Bacterial Motility Checkpoint Control. Cell Reports. 2017 Jan 24;18(4):866–77.

15. Fraebel DT, Mickalide H, Schnitkey D, Merritt J, Kuhlman TE, Kuehn S. Environment determines evolutionary trajectory in a constrained phenotypic space. Shou W, editor. eLife. 2017 Mar 27;6:e24669.

16. Morris RJ, Phan TV, Black M, Lin KC, Kevrekidis IG, Bos JA, et al. Bacterial population solitary waves can defeat rings of funnels. New J Phys. 2017 Mar;Mar(3):035002.

17. Fu X, Kato S, Long J, Mattingly HH, He C, Vural DC, et al. Spatial self-organization resolves conflicts between individuality and collective migration. Nature Communications. 2018 Jun 5;9(1):2177.

18. Cremer J, Honda T, Tang Y, Wong-Ng J, Vergassola M, Hwa T. Chemotaxis as a navigation strategy to boost range expansion. Nature. 2019 Nov;Nov(7784):658–63.

19. Liu W, Cremer J, Li D, Hwa T, Liu C. An evolutionarily stable strategy to colonize spatially extended habitats. Nature. 2019 Nov;Nov(7784):664–8.

20. Gude S, Pinçe E, Taute KM, Seinen AB, Shimizu TS, Tans SJ. Bacterial coexistence driven by motility and spatial competition. Nature. 2020 Feb;Feb(7796):588–92.

21. Phan TV, Morris R, Black ME, Do TK, Lin KC, Nagy K, et al. Bacterial Route Finding and Collective Escape in Mazes and Fractals. Phys Rev X. 2020 Jul 22;10(3):031017.

22. Bhattacharjee T, Amchin DB, Ott JA, Kratz F, Datta SS. Chemotactic migration of bacteria in porous media. Biophysical Journal. 2021 Aug 17;120(16):3483–97.

23. Bhattacharjee T, Amchin DB, Alert R, Ott JA, Datta SS. Chemotactic smoothing of collective migration. Giardina I, Walczak AM, editors. eLife. 2022 Mar 8;11:e71226.

24. Roussos ET, Condeelis JS, Patsialou A. Chemotaxis in cancer. Nat Rev Cancer. 2011 Aug;Aug(8):573–87.

25. Oppenheim JJ, Yang D. Alarmins: chemotactic activators of immune responses. Current Opinion in Immunology. 2005 Aug 1;17(4):359–65.

26. Shellard A, Mayor R. Chemotaxis during neural crest migration. Seminars in Cell & Developmental Biology. 2016 Jul 1;55:111–8.

27. Mayor R, Etienne-Manneville S. The front and rear of collective cell migration. Nat Rev Mol Cell Biol. 2016 Feb;Feb(2):97–109.

28. Patlak CS. Random walk with persistence and external bias. Bulletin of Mathematical Biophysics. 1953 Sep 1;15(3):311–38.

29. Keller EF, Segel LA. Traveling bands of chemotactic bacteria: A theoretical analysis. Journal of Theoretical Biology. 1971 Feb 1;30(2):235–48.

30. Lapidus IR, Schiller R. A model for traveling bands of chemotactic bacteria. Biophysical Journal. 1978 Apr 1;22(1):1–13.

31. Lauffenburger D, Kennedy CR, Aris R. Traveling bands of chemotactic bacteria in the context of population growth. Bulletin of Mathematical Biology. 1984 Jan 1;46(1):19–40.

32. Narla AV, Cremer J, Hwa T. A traveling-wave solution for bacterial chemotaxis with growth. Proceedings of the National Academy of Sciences. 2021 Nov 30;118(48):e2105138118.

33. Mattingly HH, Emonet T. Collective behavior and nongenetic inheritance allow bacterial populations to adapt to changing environments. Proceedings of the National Academy of Sciences. 2022 Jun 28;119(26):e2117377119.

34. Scribner TL, Segel LA, Rogers EH. A numerical study of the formation and propagation of traveling bands of chemotactic bacteria. Journal of Theoretical Biology. 1974 Jul 1;46(1):189–219.

35. Novick-Cohen A, Segel LA. A gradually slowing travelling band of chemotactic bacteria. J Math Biology. 1984 Jan 1;19(1):125–32.

36. Horstmann D. From 1970 until present: the Keller-Segel model in chemotaxis and its consequences. In 2003 [cited 2023 May 5]. Available from: https://www.semanticscholar.org/paper/From-1970-until-present%3A-the-Keller-Segel-model-in-Horstmann/332f8dd39b966aedc849956161ef644a11efa0e3

37. Wang ZA. Mathematics of traveling waves in chemotaxis --Review paper--. DCDS-B. 2012 Nov 30;18(3):601–41.

38. Seyrich M, Palugniok A, Stark H. Traveling concentration pulses of bacteria in a generalized Keller–Segel model. New J Phys. 2019 Oct 1;21(10):103001.

39. Brenner MP, Levitov LS, Budrene EO. Physical Mechanisms for Chemotactic Pattern Formation by Bacteria. Biophysical Journal. 1998 Apr 1;74(4):1677–93.

40. Marvin JS, Borghuis BG, Tian L, Cichon J, Harnett MT, Akerboom J, et al. An optimized fluorescent probe for visualizing glutamate neurotransmission. Nat Methods. 2013 Feb;Feb(2):162–70.

41. Marvin JS, Scholl B, Wilson DE, Podgorski K, Kazemipour A, Müller JA, et al. Stability, affinity, and chromatic variants of the glutamate sensor iGluSnFR. Nat Methods. 2018 Nov;Nov(11):936–9.

42. Hazel JR, Sidell BD. A method for the determination of diffusion coefficients for small molecules in aqueous solution. Analytical Biochemistry. 1987 Nov 1;166(2):335–41.

43. Ribeiro ACF, Barros MCF, Verissimo LMP, Lobo VMM, Valente AJM. Binary Diffusion Coefficients for Aqueous Solutions of l-Aspartic Acid and Its Respective Monosodium Salt. J Solution Chem. 2014 Feb 1;43(1):83–92.

44. Cremer J, Segota I, Yang C yu, Arnoldini M, Sauls JT, Zhang Z, et al. Effect of flow and peristaltic mixing on bacterial growth in a gut-like channel. PNAS. 2016 Oct 11;113(41):11414–9.

45. Monod J, Wyman J, Changeux JP. On the nature of allosteric transitions: A plausible model. Journal of Molecular Biology. 1965 May 1;12(1):88–118.

46. Neumann S, Hansen CH, Wingreen NS, Sourjik V. Differences in signalling by directly and indirectly binding ligands in bacterial chemotaxis. The EMBO Journal. 2010 Oct 20;29(20):3484–95.

47. Yang Y, M. Pollard A, Höfler C, Poschet G, Wirtz M, Hell R, et al. Relation between chemotaxis and consumption of amino acids in bacteria. Molecular Microbiology. 2015;96(6):1272–82.

48. Moore JP, Kamino K, Kottou R, Shimizu T, Emonet T. Sensory diversity and precise adaptation enable independent bet-hedging strategies for multiple signals at the same time [Internet]. bioRxiv; 2023 [cited 2023 Feb 16]. p. 2023.02.08.527720. Available from: https://www.biorxiv.org/content/10.1101/2023.02.08.527720v1

49. Mello BA, Tu Y. An allosteric model for heterogeneous receptor complexes: Understanding bacterial chemotaxis responses to multiple stimuli. PNAS. 2005 Nov 29;102(48):17354–9.

50. Colin R, Zhang R, Wilson LG. Fast, high-throughput measurement of collective behaviour in a bacterial population. Journal of The Royal Society Interface. 2014 Sep 6;11(98):20140486.

51. Dufour YS, Gillet S, Frankel NW, Weibel DB, Emonet T. Direct Correlation between Motile Behavior and Protein Abundance in Single Cells. PLOS Computational Biology. 2016 Sep 6;12(9):e1005041.

52. Mattingly HH, Kamino K, Machta BB, Emonet T. Escherichia coli chemotaxis is information limited. Nat Phys. 2021 Nov 25;1–6.

53. Mesibov R, Adler J. Chemotaxis Toward Amino Acids in Escherichia coli. Journal of Bacteriology. 1972 Oct;Oct(1):315–26.

54. Vollmer AP, Probstein RF, Gilbert R, Thorsen T. Development of an integrated microfluidic platform for dynamic oxygen sensing and delivery in a flowing medium. Lab Chip. 2005 Sep 20;5(10):1059–66.

55. Waite AJ, Frankel NW, Dufour YS, Johnston JF, Long J, Emonet T. Non-genetic diversity modulates population performance. Molecular Systems Biology. 2016 Dec 1;12(12):895.

56. Salek MM, Carrara F, Fernandez V, Guasto JS, Stocker R. Bacterial chemotaxis in a microfluidic T-maze reveals strong phenotypic heterogeneity in chemotactic sensitivity. Nature Communications. 2019 Apr 23;10(1):1877.

57. Colin R, Drescher K, Sourjik V. Chemotactic behaviour of Escherichia coli at high cell density. Nat Commun. 2019 Nov 25;10(1):5329.

58. Gurtner GC, Werner S, Barrandon Y, Longaker MT. Wound repair and regeneration. Nature. 2008 May;May(7193):314–21.

59. de Oliveira S, Rosowski EE, Huttenlocher A. Neutrophil migration in infection and wound repair: going forward in reverse. Nat Rev Immunol. 2016 Jun;Jun(6):378–91.

60. Di Talia S, Vergassola M. Waves in Embryonic Development. Annual Review of Biophysics. 2022;51(1):327–53.

61. Flemming HC, Wingender J, Szewzyk U, Steinberg P, Rice SA, Kjelleberg S. Biofilms: an emergent form of bacterial life. Nat Rev Microbiol. 2016 Sep;Sep(9):563–75.

62. Liu J, Prindle A, Humphries J, Gabalda-Sagarra M, Asally M, Lee D yeon D, et al. Metabolic co-dependence gives rise to collective oscillations within biofilms. Nature. 2015 Jul;Jul(7562):550–4.

63. Prindle A, Liu J, Asally M, Ly S, Garcia-Ojalvo J, Süel GM. Ion channels enable electrical communication in bacterial communities. Nature. 2015 Nov;Nov(7576):59–63.

64. Liu J, Martinez-Corral R, Prindle A, Lee D yeon D, Larkin J, Gabalda-Sagarra M, et al. Coupling between distant biofilms and emergence of nutrient time-sharing. Science. 2017 May 12;356(6338):638–42.

65. Petkova MD, Tkačik G, Bialek W, Wieschaus EF, Gregor T. Optimal Decoding of Cellular Identities in a Genetic Network. Cell. 2019 Feb 7;176(4):844–855.e15.

66. Díaz-Pascual F, Lempp M, Nosho K, Jeckel H, Jo JK, Neuhaus K, et al. Spatial alanine metabolism determines local growth dynamics of Escherichia coli colonies. Xavier KB, Storz G, Xavier KB, editors. eLife. 2021 Nov 9;10:e70794.

67. Jo J, Cortez KL, Cornell WC, Price-Whelan A, Dietrich LE. An orphan cbb3-type cytochrome oxidase subunit supports Pseudomonas aeruginosa biofilm growth and virulence. Donohue T, editor. eLife. 2017 Nov 21;6:e30205.

68. Douarche C, Buguin A, Salman H, Libchaber A. E. Coli and Oxygen: A Motility Transition. Phys Rev Lett. 2009 May 12;102(19):198101.

69. Aggarwal A, Liu R, Chen Y, Ralowicz AJ, Bergerson SJ, Tomaska F, et al. Glutamate indicators with improved activation kinetics and localization for imaging synaptic transmission. Nat Methods. 2023 May 4;1–10.

70. Studier FW. Protein production by auto-induction in high-density shaking cultures. Protein Expression and Purification. 2005 May 1;41(1):207–34.

71. Crowe J, Dobeli H, Gentz R, Hochuli E, Stiiber D, Henco K. 6xffis-Ni-NTA Chromatography as a Superior Technique in Recombinant Protein Expression/Purification. In: Harwood AJ, editor. Protocols for Gene Analysis [Internet]. Totowa, NJ: Humana Press; 1994 [cited 2023 May 8]. p. 371–87. (Methods in Molecular Biology). Available from: https://doi.org/10.1385/0-89603-258-2:371

72. Soupene E, van Heeswijk WC, Plumbridge J, Stewart V, Bertenthal D, Lee H, et al. Physiological Studies of Escherichia coli Strain MG1655: Growth Defects and Apparent Cross-Regulation of Gene Expression. Journal of Bacteriology. 2003 Sep 15;185(18):5611–26.

73. Lyons E, Freeling M, Kustu S, Inwood W. Using Genomic Sequencing for Classical Genetics in E. coli K12. PLOS ONE. 2011 Feb 25;6(2):e16717.

74. You C, Okano H, Hui S, Zhang Z, Kim M, Gunderson CW, et al. Coordination of bacterial proteome with metabolism by cyclic AMP signalling. Nature. 2013 Aug;Aug(7462):301–6.

75. Hui S, Silverman JM, Chen SS, Erickson DW, Basan M, Wang J, et al. Quantitative proteomic analysis reveals a simple strategy of global resource allocation in bacteria. Molecular Systems Biology. 2015 Feb;Feb(2):784.

76. Brown SD, Jun S. Complete Genome Sequence of Escherichia coli NCM3722. Genome Announcements. 2015 Aug 6;3(4):e00879–15.

77. Basan M, Hui S, Okano H, Zhang Z, Shen Y, Williamson JR, et al. Overflow metabolism in Escherichia coli results from efficient proteome allocation. Nature. 2015 Dec;Dec(7580):99–104.

78. Basan M, Honda T, Christodoulou D, Hörl M, Chang YF, Leoncini E, et al. A universal trade-off between growth and lag in fluctuating environments. Nature. 2020 Aug;Aug(7821):470–4.

79. Shen Y, Chen Y, Wu J, Shaner NC, Campbell RE. Engineering of mCherry variants with long Stokes shift, red-shifted fluorescence, and low cytotoxicity. PLOS ONE. 2017 Feb 27;12(2):e0171257.

80. Edelstein A, Amodaj N, Hoover K, Vale R, Stuurman N. Computer Control of Microscopes Using μManager. Current Protocols in Molecular Biology. 2010;92(1):14.20.1–14.20.17.

