## Supporting Information for "Direct measurement of dynamic attractant gradients reveals breakdown of the Patlak-Keller-Segel chemotaxis model"

**This PDF file includes:**

Supporting text
Figures S1 and S2
SI References

### Supporting Information Text

#### Local drop in oxygen concentration is expected to be proportional to bacterial density.

We model dissolved oxygen concentration $O$ with the following dynamics, averaged over the width of the channel:

$$\partial_t O = D_O\left(\partial_{xx} O + \partial_{yy} O\right) - k_o \frac{O}{O + K_O} B(x, y)$$

where $x$ is the direction of the long channel, $y$ is the vertical position in the channel, $B$ is bacterial density, $D_O$ is the diffusivity of oxygen in water, $k_O$ is the maximum consumption rate of oxygen by bacteria, and $K_O$ is the Monod constant of oxygen consumption. Because the channel depth $L = 100\ \mu\text{m}$ is shallow compared to its width (1.2 mm), we can neglect transfer of oxygen into the channel from the side walls.

This PDE is paired with boundary conditions:

$$O(x, y = 0) = O_{out}.$$

$$\left[\partial_y O(x, y)\right]_{y=L} = 0$$

The first boundary condition fixes the oxygen concentration at the water-PDMS interface to $O_{out}$. The second condition reflects the fact that the glass slide is impermeable to oxygen.

Assuming that $K_O$ is small compared to the minimum oxygen concentration, that variations in cell density $B$ in the $y$ direction are negligible, and that the cells form a quasi-stationary traveling wave with moving reference frame $z = x - c\,t$, we get:

$$-c\,\partial_z O = D_O\left(\partial_{zz} O + \partial_{yy} O\right) - k_o\, B(z).$$

Since the wave is much wider than the depth of the channel, variations in oxygen along the $y$-direction are much larger than those in the $z$-direction:

$$\partial_{yy} O - \frac{k_o}{D_O} B(z) = 0$$

Integrating once:

$$\partial_y O - \frac{k_o}{D_O} B(z)\, y + a_1 = 0$$

Applying the boundary condition at $y = L$:

$$-\frac{k_o}{D_O} B(z)\, L\ + a_1 = 0$$

$$a_1 = \frac{k_o}{D_O} B(z)\, L$$

$$\partial_y O - \frac{k_o}{D_O} B(z)\,(y - L) = 0$$

Integrating again:

$$O(z, y) - \frac{k_o}{D_O} B(z)\left(\frac{y^2}{2} - L\,y\right) + a_2 = 0$$

Applying the boundary condition at $y = 0$:

$$O_{out} + a_2 = 0$$

$$a_2 = -O_{out}.$$

Together:

$$O(z,y) - \frac{k_o}{D_O} B(z)\, y\left(\frac{y}{2} - L\right) - O_{out} = 0$$

$$O_{out} - O(z,y) = \frac{k_o}{D_O} B(z)\, y\left(L - \frac{y}{2}\right).$$

Averaging the drop in oxygen over the channel depth:

$$\langle \Delta O \rangle_y(z) = \langle O_{out} - O(z,y) \rangle_y = \frac{1}{L}\int_0^L \frac{k_o}{D_O} B(z)\, y\left(L - \frac{y}{2}\right) dy$$

$$= \frac{1}{L}\frac{k_o}{D_O} B(z)\left(L\frac{L^2}{2} - \frac{L^3}{6}\right)$$

$$= \frac{L^2}{3}\frac{k_o}{D_O} B(z).$$

Thus, we expect that the local drop in oxygen concentration at position $z$ in the wave scales like the channel depth squared, $L^2$, and is proportional to the local cell density $B(z)$.

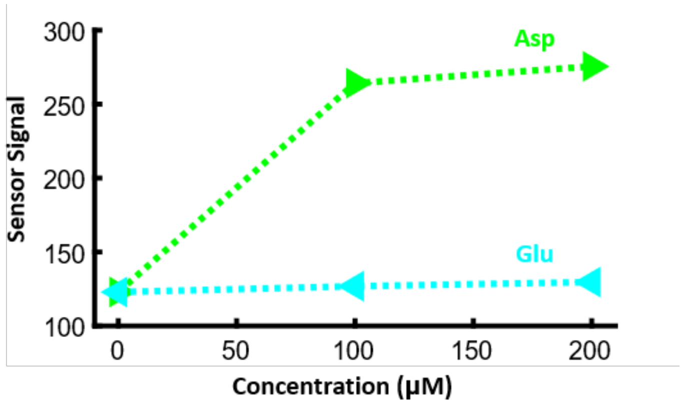


**Fig. S1. iAspSnFR is specific for aspartate and insensitive to glutamate.** Sensor fluorescence emission response to glutamate (Glu, cyan) is negligible compared to the response to aspartate (Asp, green). Concentration values are $\mathbf{0}$, $\mathbf{100}$ and $\mathbf{200\ \mu M}$ for both amino acids. Data points are averages over an entire sweep of images along the microfluidic device. The same device was used for all Asp concentrations.

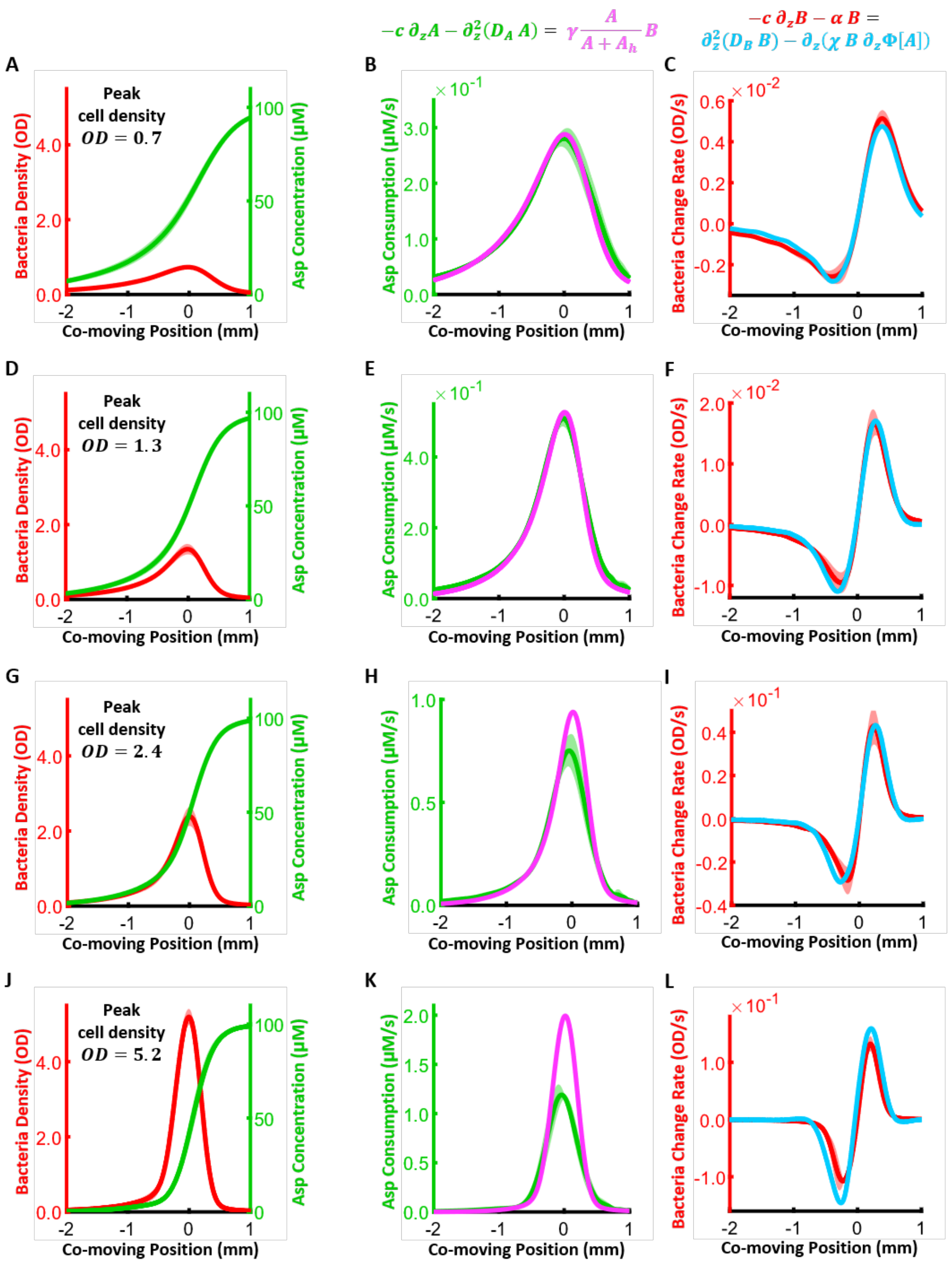


**Fig. S2. The standard model for chemotaxis, the Patlak-Keller-Segel (PKS) model, describes the bacterial wave at low cell densities but breaks down at high densities**. **(A)** The quasi-steady state profiles in the co-moving frame, of bacteria density (red) and attractant Asp concentration (green), during low cell density migration (OD = 0.7). **(B)** Fitting the Asp dynamics of the PKS model to experimental data (above). $\boldsymbol{A}$ and $\boldsymbol{B}$ are the Asp concentration and the bacteria density respectively, $\boldsymbol{c}$ is the wave speed, $\boldsymbol{D_A}$ is the molecular diffusion of Asp, $\boldsymbol{\gamma}$ is the maximum consumption rate, $\boldsymbol{A_h}$ is the half-max of the consumption rate, and $\boldsymbol{z = x - c\,t}$ is the coordinate in the co-moving frame ($\boldsymbol{z = 0}$ is set to the peak bacteria density). Green: LHS of the Asp dynamics equation with $\boldsymbol{A}$ and $\boldsymbol{c = 4.4 \pm 0.1\ \mu m/s}$ measured and $\boldsymbol{D_A = 800\ \mu m^2/s}$ (1–5). Magenta: RHS of the Asp dynamics equation with $\boldsymbol{B}$ measured, and $\boldsymbol{\gamma = 0.45 \pm 0.04\ \mu M/OD/s}$ and $\boldsymbol{A_h = 7.6 \pm 3.7\ \mu M}$ fit to match the LHS. **(C)** Fitting the bacterial dynamics of the PKS model to experimental data (above). $\boldsymbol{\Phi[A] = \ln\left(\frac{1+A/K_i}{1+A/K_a}\right)}$ is the cells' perceived signal ($\boldsymbol{K_a \gg A}$ and $\boldsymbol{K_i = 1\ \mu M}$ (6–9)), $\boldsymbol{D_B}$ is the effective bacterial diffusivity, $\boldsymbol{\chi}$ is the chemotactic coefficient, and $\boldsymbol{\alpha = 0.4/hr}$ is the measured growth rate (Methods). Red: LHS of the bacteria dynamics equation with $\boldsymbol{\alpha}$, $\boldsymbol{c}$ and $\boldsymbol{B}$ measured. Cyan: RHS of the bacteria dynamics equation with $\boldsymbol{D_B = 400 \pm 100\ \mu m^2/s}$ and $\boldsymbol{\chi = 3300 \pm 230\ \mu m^2/s}$ fit to match the LHS. **(D,E,F), (G,H,I), (J,K,L)** are the same as **(A,B,C)** for high cell density waves. Velocities are $\boldsymbol{c} = 5.5 \pm 0.1\ \boldsymbol{\mu m/s}$ (OD = 1.3), $\boldsymbol{c = 6.4 \pm 0.2\ \mu m/s}$ (OD = 2.4) and $\boldsymbol{c = 7.7 \pm 0.2\ \mu m/s}$ (OD = 5.2). **(E,F), (H,I), (K,L)** magenta and cyan lines are prediction using the same parameter values for $\boldsymbol{\gamma}$, $\boldsymbol{A_h}$, $\boldsymbol{D_B}$, $\boldsymbol{\chi}$ as in **(B,C)**. The shading represents the standard deviation across replicates. Replicate $\boldsymbol{N = 6}$ for **(A,B,C)**, $\boldsymbol{N = 9}$ for **(D,E,F)**, $\boldsymbol{N = 5}$ for **(G,H,I)**, and $\boldsymbol{N = 4}$ for **(J,K,L)**.

**SI References**


1. Fu X, Kato S, Long J, Mattingly HH, He C, Vural DC, et al. Spatial self-organization resolves conflicts between individuality and collective migration. Nature Communications. 2018 Jun 5;9(1):2177.

2. Cremer J, Honda T, Tang Y, Wong-Ng J, Vergassola M, Hwa T. Chemotaxis as a navigation strategy to boost range expansion. Nature. 2019 Nov;575(7784):658–63.

3. Hazel JR, Sidell BD. A method for the determination of diffusion coefficients for small molecules in aqueous solution. Analytical Biochemistry. 1987 Nov 1;166(2):335–41.

4. Ribeiro ACF, Barros MCF, Verissimo LMP, Lobo VMM, Valente AJM. Binary Diffusion Coefficients for Aqueous Solutions of l-Aspartic Acid and Its Respective Monosodium Salt. J Solution Chem. 2014 Feb 1;43(1):83–92.

5. Cremer J, Segota I, Yang C yu, Arnoldini M, Sauls JT, Zhang Z, et al. Effect of flow and peristaltic mixing on bacterial growth in a gut-like channel. PNAS. 2016 Oct 11;113(41):11414–9.

6. Neumann S, Hansen CH, Wingreen NS, Sourjik V. Differences in signalling by directly and indirectly binding ligands in bacterial chemotaxis. The EMBO Journal. 2010 Oct 20;29(20):3484–95.

7. Yang Y, M. Pollard A, Höfler C, Poschet G, Wirtz M, Hell R, et al. Relation between chemotaxis and consumption of amino acids in bacteria. Molecular Microbiology. 2015;96(6):1272–82.

8. Moore JP, Kamino K, Kottou R, Shimizu T, Emonet T. Sensory diversity and precise adaptation enable independent bet-hedging strategies for multiple signals at the same time [Internet]. bioRxiv; 2023 [cited 2023 Feb 16]. p. 2023.02.08.527720. Available from: https://www.biorxiv.org/content/10.1101/2023.02.08.527720v1

9. Mello BA, Tu Y. An allosteric model for heterogeneous receptor complexes: Understanding bacterial chemotaxis responses to multiple stimuli. PNAS. 2005 Nov 29;102(48):17354–9.
